# Brain structure-function coupling – relationship with language lateralisation

**DOI:** 10.1101/2025.09.05.674563

**Authors:** Ieva Andrulyte, Laure Zago, Gael Jobard, Herve Lemaitre, Peter N Taylor, Francois Rheault, Marc Joliot, Laurent Petit, Simon S Keller

## Abstract

Language is one of the most extensively studied lateralised cognitive functions in the human brain, predominantly relying on the left hemisphere in most individuals. However, the mechanisms by which a stable white matter architecture underpins individual language functions remain unclear. Previous studies have employed structural connectivity (SC) and functional connectivity (FC) coupling for individual fingerprinting and task decoding, suggesting that variability in brain entropy may serve as a distinguishing characteristic for language lateralisation. We examined a large cohort of healthy adults (n = 285) to investigate SC-FC coupling and identify markers distinguishing different language laterality groups. Functional connectivity was measured using resting-state fMRI (rsfMRI) time-series data, while structural connectivity was determined via probabilistic fibre tractography. SC-FC coupling was investigated using the SENSAAS language atlas and defined as the Pearson correlation between non-zero elements of regional structural and functional connectivity profiles. Group differences were assessed using the PALM toolbox in FSL. Our findings revealed that increased SC-FC coupling in the left precentral sulcus was associated with typical language lateralisation, while increased coupling in the right middle temporal gyrus and left anterior insula was observed in individuals with atypical language lateralisation (pFDR < 0.05). Non-lateralised individuals exhibited increased coupling in the left anterior insula compared to lateralised (pFDR<0.05). SC-FC coupling offers a promising framework to uncover functional and anatomical differences among individuals with varying language lateralisation. This regional specificity indicates that typical, atypical, and non-lateralised profiles rely on different structural-functional alignments, likely reflecting the recruitment of alternative pathways for language processing.

## Introduction

Hemispheric language dominance refers to the brain’s interhemispheric asymmetry in language function, with around 90% of healthy individuals relying primarily on the left hemisphere, while the remaining 10% exhibit atypical patterns, characterised by bilateral involvement or primary reliance on the right hemisphere (Andrulyte et al., 2024; Mazoyer et al., 2014). The study of language lateralisation has begun to adopt a connectomics framework (Andrulyte, Zago, et al., 2025; Labache et al., 2020), which conceptualises the brain as an interconnected network of structural and functional connections supporting communication, information processing, and cognitive integration (Hagmann et al., 2008). This framework encompasses structural connectivity (SC) – the anatomical wiring reconstructed through diffusion MRI and tractography (Basser et al., 1994; Maier-Hein et al., 2017; Turner et al., 1990) – and functional connectivity (FC) – the temporal synchronisation of neural activity between brain regions, measured via resting-state fMRI (rsfMRI) (Biswal et al., 1995). SC captures stable anatomical pathways, while FC reflects dynamic neural interactions that support cognitive processing and neural communication (Avena-Koenigsberger et al., 2018; Betzel, 2020; Sotiropoulos & Zalesky, 2019; Tao et al., 2015).

Advances in rsfMRI and diffusion MRI (dMRI) analysis methods have enabled network-based approaches for the investigation of the neural basis of language lateralisation. In healthy individuals, rsfMRI studies have found that atypical lateralisation is associated with greater interhemispheric connectivity (Labache et al., 2020) and weaker intra-hemispheric integration (Ibrahim et al., 2015). dMRI analyses further support these findings, showing increased fractional anisotropy (FA) in interhemispheric temporal regions (Andrulyte et al., 2024; Andrulyte, Zago, et al., 2025) and higher nodal degree in the posterior middle temporal gyrus and posterior cingulate cortex in atypically lateralised people (Zahnert et al., 2023).

While rsfMRI and dMRI have independently contributed to our understanding of the connectome, their integration offers a more comprehensive framework for studying language lateralisation. SC-FC coupling – the relationship between the brain’s structural architecture and functional activity – provides valuable insights into how neural circuits support cognitive processes (Skudlarski et al., 2008; Suárez et al., 2020). Hebb’s principle, “neurons that fire together wire together,” highlights that repeated co-activation of neurons strengthens synaptic connections, shaping structural networks that optimise neural processing (Hebb, 1949; Straathof et al., 2019). Although structural connectivity explains 50–60% of the variance in functional connectivity, deviations between the two (decoupling) are particularly prominent in higher-order brain regions such as the default mode network, which supports integrative cognitive functions (Baum et al., 2020; Preti & Van De Ville, 2019; Sarwar et al., 2021). This hierarchical organisation reflects an evolutionary specialisation: tightly coupled unimodal regions support efficient sensory processing, while more flexible coupling in transmodal regions supports integrative and adaptive functions (Huntenburg et al., 2017; Margulies et al., 2016; Yeo et al., 2016).

Coupling between structural and functional networks is thought to underpin individual differences in cognitive flexibility and specialisation (Medaglia et al., 2017). For example, weaker coupling in transmodal regions has been linked to cognitive flexibility, promoting regional specialisation, which allows the brain to support unique behavioural repertoires (Preti & Van De Ville, 2019). The critical brain hypothesis, which suggests the brain operates near a balance of order and chaos to optimise adaptability, aligns with these principles (Shi et al., 2021). While originally grounded in electroencephalography (EEG) studies, it may extend to structure-function coupling, where maintaining such a balance could support efficient information processing. Disruptions to this balance, observed in conditions such as epilepsy and schizophrenia, impair cognition by destabilising the relationship between structural and functional connectivity (Fornito & Bullmore, 2015; Wu et al., 2020; Z. Zhang et al., 2011).

Although coupling between structural and functional networks has proven useful in profiling cognitive traits, its application to language lateralisation has been limited but promising. Effective connectome analyses, which infer causal relationships between structural pathways and functional interactions, have revealed distinct language-related networks that connect sensory and self-referential systems with syntactic and motor planning regions, highlighting the structural-functional basis of language processes (Rolls et al., 2022). One study, focusing specifically on language lateralisation, employed diffusion map embedding to derive individual functional connectomes. It found that left-lateralised individuals exhibit increased resting-state connectivity in key regions such as the left inferior frontal gyrus, left inferior parietal lobule, and left middle temporal gyrus (Langs et al., 2016). These studies highlight the promise of integrating rsfMRI and dMRI for understanding language lateralisation, yet no research to date has explicitly examined SC-FC coupling as a potential marker for language dominance.

This study aims to bridge existing gaps by investigating the relationship between connectivity strength in diffusion MRI and rsfMRI modalities and their coupling in relation to language lateralisation. Specifically, we have two key objectives: (1) to examine rsfMRI and diffusion MRI FA connectivity strength at the level of individual nodes to identify associations with language lateralisation, and (2) to assess macroscale SC-FC coupling strength and its relationship with functional language lateralisation. To achieve this, we leverage the BIL&GIN dataset, which provides well-characterised classifications of individuals with typical and atypical language lateralisation (Mazoyer et al., 2016). By analysing structure-function coupling in these groups, this study aimed to evaluate its potential as a marker for language dominance. We hypothesise that distinct coupling patterns will emerge across groups, with left-lateralised individuals showing increased coupling in classical language areas, such as the inferior frontal gyrus and middle temporal gyrus in the dominant hemisphere, and atypically lateralised individuals exhibiting reduced coupling.

## Methods

### Data acquisition

Structural and functional neuroimaging data were sourced from the BIL&GIN database, specifically designed to investigate the structural and functional neural correlates of brain lateralisation (Mazoyer et al., 2016). This study utilised data from 285 healthy individuals (aged 18–57 years; 48% female), including individual measures derived from diffusion MRI and resting-state functional MRI (rsfMRI) time-series.

Imaging data were acquired using a Philips Achieva 3-Tesla MRI scanner. The structural MRI protocol included a localiser scan and a high-resolution 3D T1-weighted sequence. The T1-weighted imaging parameters were as follows: repetition time (TR) = 20 ms, echo time (TE) = 4.6 ms, flip angle = 10°, inversion time = 800 ms, turbo field echo factor = 65, SENSE factor = 2, field of view (FOV) = 256 × 256 × 180 mm^3^, and isotropic voxel size = 1 mm^3^. Functional data were acquired using T2*-weighted echo-planar imaging (EPI) with TR = 2 s, TE = 35 ms, flip angle = 80°, and 31 axial slices, with an isotropic voxel size of 3.75 × 3.75 × 3.75 mm^3^. The FOV was consistent with the T2*-FFE acquisition. Functional imaging for a sentence production task (Mazoyer et al., 2016) included three runs, each comprising 192 T2*-weighted volumes. For rsfMRI, participants underwent an 8-minute scan with instructions to keep their eyes closed, stay relaxed, minimise movement, remain awake, and let thoughts flow freely.

Diffusion-weighted imaging (DWI) employed a single-shot spin-echo EPI sequence. The protocol included a b0 image (b = 0 s/mm^2^) and 21 non-collinear diffusion gradient directions (b = 1000 s/mm^2^), each acquired twice with reversed gradient polarities. Gradient directions were evenly distributed across a half-sphere, with complementary directions on the opposing half-sphere. To enhance signal-to-noise ratio, the acquisition was repeated, resulting in a total scan time of 15 minutes and 30 seconds. Seventy axial slices were collected, aligned parallel to the anterior commissure–posterior commissure (AC–PC) plane, spanning from the cerebellum base to the vertex. The imaging parameters included TR = 8500 ms, TE = 81 ms, flip angle = 90°, SENSE reduction factor = 2.5, acquisition matrix = 112 × 112, and an isotropic voxel size of 2 × 2 × 2 mm^3^.

### Language lateralisation

To assess language laterality, we utilised the sentence production task from the BIL&GIN dataset (Mazoyer et al., 2016) during a task-based fMRI session, employing a word-list production contrast. For the sentence production task, participants were presented with stimuli derived from the classic French comic strip series Le Petit Nicolas. After observing a “Le petit Nicolas” line drawing, participants were instructed to covertly generate a sentence structured as follows: a subject and complement, followed by a verb describing the action, and concluding with an additional complement specifying place or manner. In contrast, during the reference word production task, scrambled drawings served as stimuli, and participants were instructed to covertly generate a list of the months of the year. Further details on these tasks can be found in Mazoyer et al. (2016).

Language production laterality indices (LIs) were computed using the LI toolbox (Wilke & Schmithorst, 2006). The analysis was conducted within grey and white matter anatomical template masks, excluding the cerebellum, to maintain consistency with fMRI data normalisation protocols. The LI toolbox generated 10,000 potential LI combinations, and the central 50% were retained to mitigate the influence of statistical outliers. Final HFLI values, ranging from −100 (indicating exclusive right-hemisphere activation) to +100 (exclusive left-hemisphere activation), were calculated by weighting the central 50% of LIs based on their corresponding thresholds. To categorise individuals as lateralised, we used an LI threshold of 20, classifying individuals with |LI| > 20 as lateralised and those with |LI| ≤ 20 as non-lateralised (i.e. bilateral).

To distinguish between atypically and typically lateralised individuals, we applied a threshold, defining individuals with LI > 20 as “typically” lateralised and those with lower values as “atypically” lateralised (Chang et al., 2017). Unlike prior studies (Andrulyte, Zago, et al., 2025), where atypical individuals were divided into two subgroups (e.g., right-lateralised and non-lateralised), we combined these groups into a single atypical category here to enhance interpretability and statistical power. In this study, we examined the effects of lateralisation versus bilaterality and typical versus atypical lateralisation separately, aiming to identify distinct components contributing to these differences.

### Functional MRI data processing

Both resting-state and task-related fMRI data were processed using the global linear modelling software Statistical Parametric Mapping (SPM5; www.fil.ion.ucl.ac.uk/spm/) alongside customised MATLAB routines. Functional data from the three slow fMRI runs (gradient echo planar imaging, repetition time [TR] = 2.0 s, voxel size = 3.75 × 3.75 × 3.75 mm^3^; acquired on a 3T Philips Intera Achieva scanner) underwent slice-timing and motion correction. Movement artefacts were mitigated by regressing the six motion parameters (three translations and three rotations) from each voxel’s T2*-EPI time series. The corrected T2*-EPI scans were rigidly co-registered to the participants’ structural T2*-FFE images.

The T1-weighted (T1w) volumes were segmented into grey matter, white matter, and cerebrospinal fluid. Spatial normalisation was performed by integrating the transformation matrix from T2*-FFE to T1w images with the stereotaxic normalisation matrix derived from T1w images. This process enabled the warping of T2*-EPI scans into standard stereotaxic space (2 × 2 × 2 mm^3^ voxel resolution) using a single trilinear interpolation. Finally, voxel time series were high-pass filtered with a 159-second cut-off to remove low-frequency drifts (Joliot et al., 2016).

### Resting-state fMRI connectivity mapping

Functional connectivity was assessed by extracting resting-state fMRI time-series data from 64 supramodal sentence-related regions defined by the SENSAAS language atlas, comprising 32 left-lateralised ROIs and their homotopic counterparts in the right hemisphere (Labache et al., 2019) (Figure 1). Pearson correlation coefficients were then computed between these regions to generate a correlation matrix capturing interregional functional connectivity (Sarwar et al., 2021). To improve clarity and facilitate interpretation, negative correlations within each connectivity matrix were replaced with zero values (Luppi & Stamatakis, 2021).

**Figure 1.**
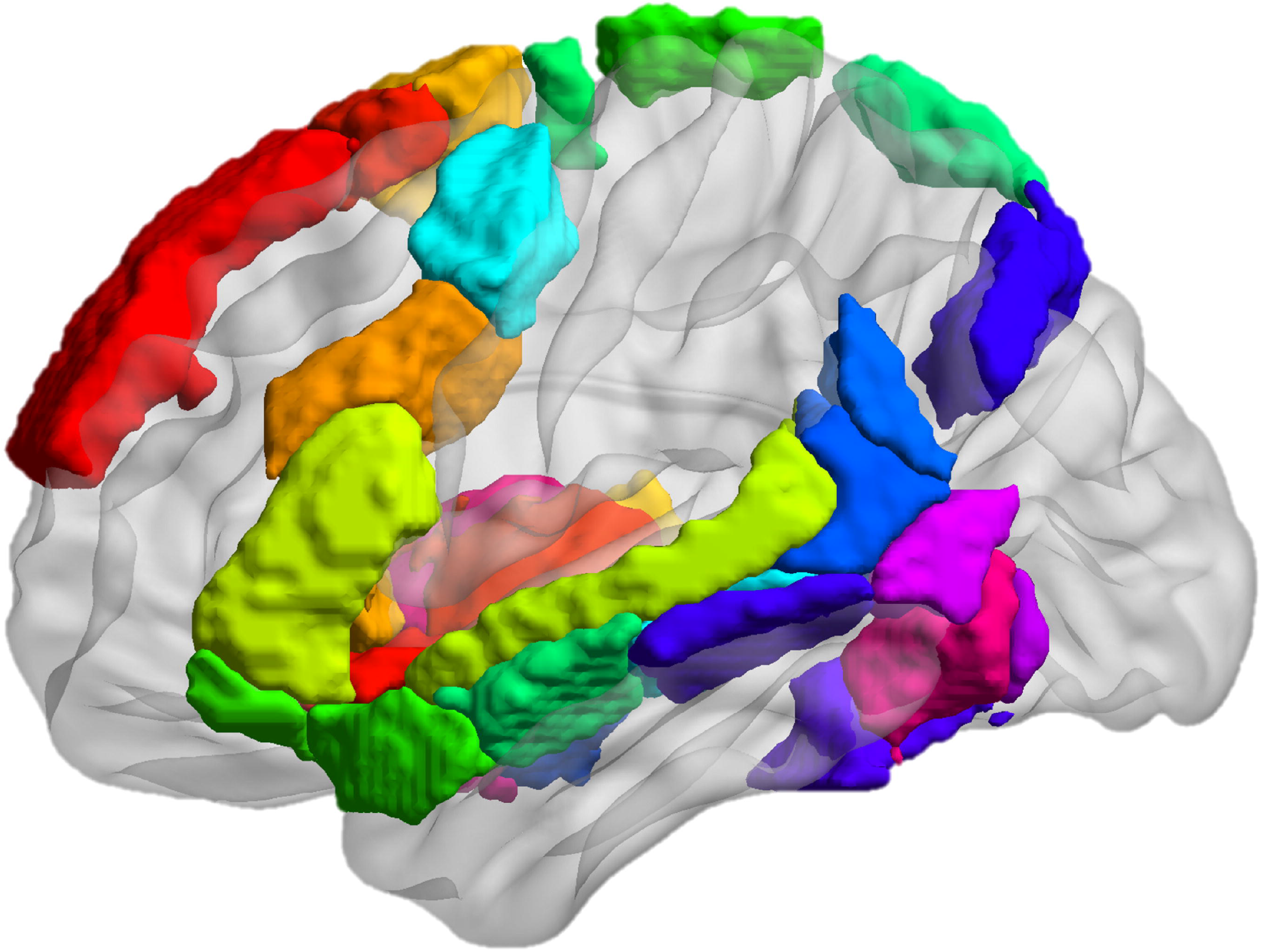
Brain regions included in the analysis. Laterality indices (LIs) were calculated across the whole brain using the AICHA atlas (Joliot et al., 2015). The SENSAAS atlas, developed for the BIL&GIN dataset, was then used to define regions of interest implicated in sentence production (Labache et al., 2019).

### Diffusion MRI data processing

Probabilistic tractography was performed using TractoFlow, an automated diffusion MRI processing pipeline. TractoFlow integrates two tracking algorithms – classical local tracking and Particle Filter Tracking (PFT) (Girard et al., 2014) – to generate whole-brain tractograms. Diffusion MRI data were acquired using a spin-echo echo-planar imaging sequence with a b0 map and 84 diffusion-weighted images (b = 1000 s/mm^2^), which were averaged in sets of four to yield 21 unique non-collinear diffusion directions. The diffusion signal was modelled using spherical harmonics of order 6 (Descoteaux et al., 2009). Whole-brain seeding was conducted, with PFT employing 10 seeds per voxel and local tracking using 5 seeds per voxel. The default TractoFlow parameters were used, including a step size of 0.5 mm and a maximum angular deviation of 20°between steps.

To ensure anatomical consistency, tractograms were normalised to MNI space using ANTs (Avants et al., 2009). The transformation parameters, including both affine and nonlinear components derived from the registration of T1-weighted images to the MNI152 2009 standard-space template (1 mm isotropic resolution; Legarreta et al., 2021), were applied to warp the whole-brain tractograms.

To refine the tractograms, false-positive tracts were removed using the ExTractorFlow pipeline, which applies rule-based filtering grounded in neuroanatomical organisational principles. The method classifies streamlines as anatomically implausible if they violate known white matter architecture, such as forming 360° loops, terminating along the ventricles, fragmenting within deep white matter structures, or being shorter than 20 mm. This ensures that only streamlines following known pathways – association, commissural, and projection fibres – are retained. Further methodological details can be found in Petit et al. (2023). Thus, this approach enhances reproducibility and precision compared to conventional diffusion MRI techniques, enabling more accurate identification of anatomically plausible connections through region-of-interest filtering.

### Diffusion MRI connectivity mapping

Structural connectivity was assessed using the same AICHA parcellation as in the rsfMRI analysis (Joliot et al., 2015), defining 384 brain regions (192 per hemisphere). Whole-brain tractograms were segmented into connectivity matrices using Scilpy toolbox (https://github.com/scilus/scilpy), where nodes corresponded to the parcellated brain regions and edges represented the white matter tracts linking them. Edges were weighted by fractional anisotropy (FA) values, which were chosen over streamline count due to its reduced susceptibility to biases inherent in probabilistic tractography (F. Zhang et al., 2022). While FA is sensitive to crossing fibres and other microstructural complexities (Jeurissen et al., 2013; Tournier et al., 2011), it remains widely used for structural network analyses (Lebel et al., 2019) and is a primary metric of interest in investigations of language lateralisation (Andrulyte, Demirkan, et al., 2025). The 384 × 384 connectivity matrices were downsampled to 64 × 64 by extracting 32 left-lateralised supramodal sentence-processing regions and their homotopic counterparts, as defined by the SENSAAS language atlas (Labache et al., 2019) (Figure 1).

### Connectivity strengths for structural and functional connectivity and group comparisons

We calculated the average connectivity for each node in the SENSAAS atlas (Labache et al., 2019) in each subject using connectivity matrices derived separately from FA-based structural connectivity and rsfMRI functional connectivity. For readability, we use the term “connectivity strength”; however, in diffusion MRI, FA reflects diffusion properties rather than direct measures of connection efficacy or information transfer and should be interpreted accordingly (Figley et al., 2022).

For each modality, group comparisons were conducted using independent t-tests to assess differences between typically and atypically lateralised individuals, as well as lateralised versus non-lateralised groups. Effect sizes (Cohen’s *d*) were computed to quantify the magnitude of these differences, with values of 0.2, 0.5, and 0.8 corresponding to small (∼85% population overlap), medium (∼67% overlap), and large (∼53% overlap) effects, respectively (Cohen, 1977). P-values were corrected for multiple comparisons using the false discovery rate (FDR), and nodes showing significant effects were identified.

### Structure-function coupling

Structure–function coupling was computed using the 64 regions defined by the SENSAAS atlas (Labache et al., 2019). For each node, coupling was defined as the Pearson correlation between the non-zero elements of its regional structural connectivity profile (measured using fractional anisotropy, FA) and its functional connectivity profile (measured using resting-state fMRI correlation coefficients) (Baum et al., 2020) (Figure 2). Regional connectivity profiles for each participant were represented as vectors of connectivity strength from a single network node to all other nodes, excluding diagonal elements (Zarkali et al., 2021). This approach yielded a single coupling value per node, reflecting the relationship between its structural and functional connectivity patterns. Fisher’s Z-transformation was applied to the resulting correlation coefficients to normalise the data for statistical analysis.

**Figure 2.**
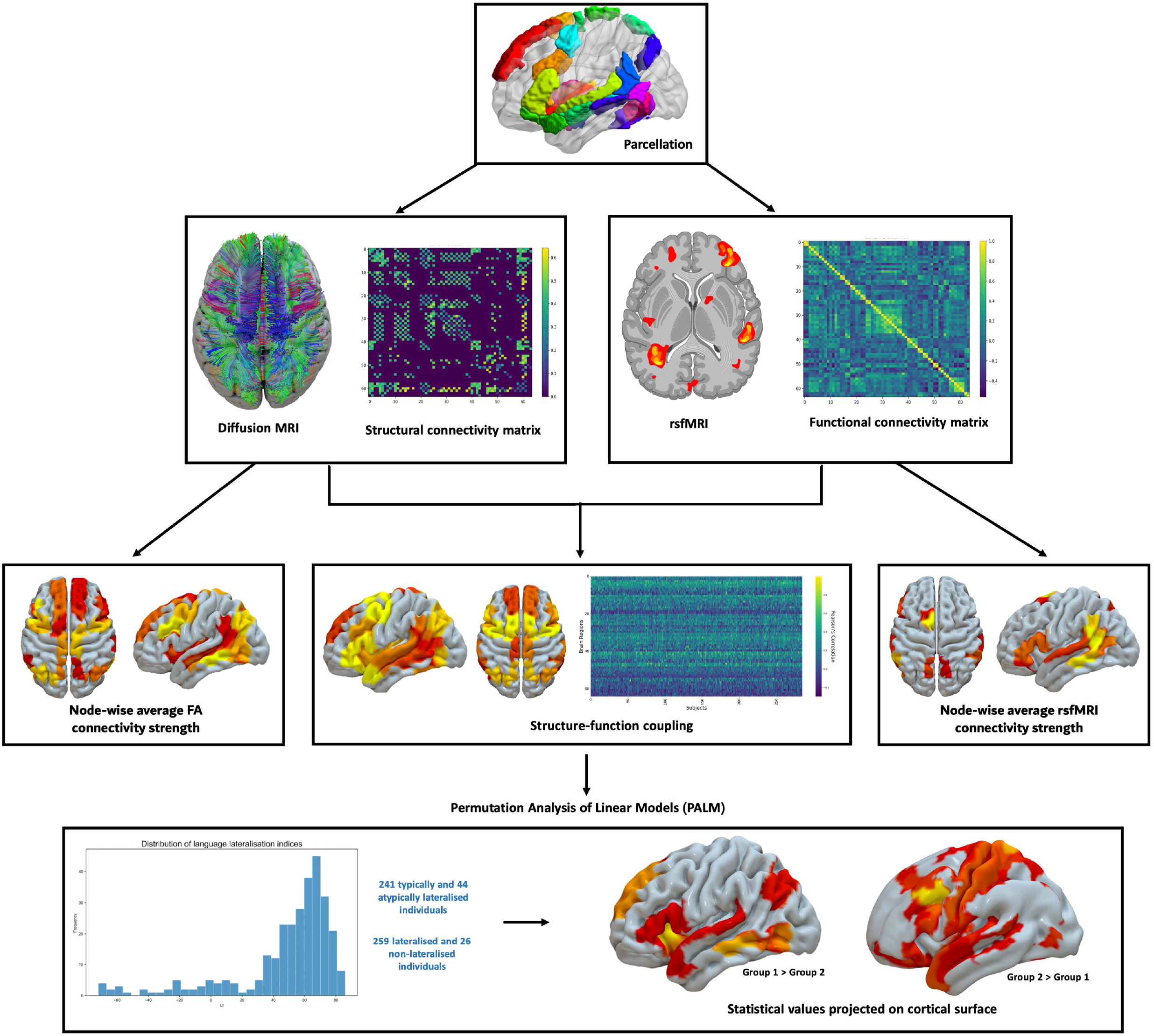
Pipeline for computing structure-function coupling. This figure illustrates the methodological steps for assessing structure-function coupling within the brain. The brain is first divided into distinct regions of interest (nodes) using a predefined atlas. Diffusion MRI data is processed to generate a structural connectivity matrix. This matrix quantifies the strength of white matter connections between each pair of nodes, using fractional anisotropy (FA) as a measure of connectivity. Resting-state fMRI (rsfMRI) data is used to create a functional connectivity matrix, representing the temporal correlations in neuronal activity between each pair of nodes, calculated using Pearson’s correlation. For each node, average FA connectivity strength is computed as the mean FA value of its structural connections, and average rsfMRI connectivity strength is computed as the mean functional connectivity value of its connections. Structure-function coupling is computed by correlating corresponding entries in the structural and functional connectivity matrices using Pearson’s correlation, yielding a coupling measure for each node. This results in a measure of structure-function coupling for each region. Finally, differences in structure-function coupling between groups are statistically assessed using permutation analysis for linear models (PALM) implemented in FSL with 10,000 permutations. This analysis identifies regions where the relationship between structural and functional connectivity differs significantly between the groups.

To visualise global structure-function coupling, Z-transformed coupling scores were averaged across participants for each node and mapped onto the cortical brain surface. Group comparisons were conducted using the PALM (Permutation Analysis of Linear Models) toolbox in FSL (Winkler et al., 2014), testing two contrasts: (1) bilateral vs lateralised language representation and (2) typical vs atypical language lateralisation. False discovery rate correction (FDR, *p* < 0.05) was applied across the 64 regions defined by the SENSAAS atlas to control for multiple comparisons. Covariates of no interest, including sex, age, and handedness, were incorporated into the model. Group differences were assessed using 10,000 permutations.

### Posthoc analyses

To further investigate the relationship between structural and functional connectivity in language lateralisation, we conducted post-hoc analyses focusing exclusively on edge-level coupling for connections involving significant regions. Specifically, we used MATLAB (version R2023a) to extract edge-level connectivity values from these regions to all other brain areas and quantified structure-function coupling by multiplying resting-state functional MRI (rsfMRI) connectivity values with diffusion MRI-based fractional anisotropy (FA) for each connection. As in our main analyses, negative functional connectivity values were thresholded at zero before multiplication, ensuring that only positive rsfMRI connectivity values were used in computing coupling metrics.

## Results

### Participant characteristics

Out of 285 healthy participants, 241 were typically lateralised (mean age 25.5 ± 6.17 years, 49% female), while 44 were atypically lateralised (mean age 27.0 ± 7.86 years, 46% female). In terms of bilateral versus lateralised grouping, 259 participants were lateralised (right- or left-hemisphere lateralised; mean age 25.6 ± 6.21 years, 50% female), and 26 had bilateral language representation (mean age 27.2 ± 8.63 years, 31% female). The overall mean age of participants was 25.7 years (SD ± 6.5), with no significant age differences observed between the typical and atypical groups or between the lateralised and bilateral groups (t-test, p > 0.05).

Female participants comprised 48% of the total sample (n = 138), with no significant differences in gender distribution across the lateralisation groups (chi-square, p > 0.05). Handedness was evenly distributed, with 48% of participants being right-handed and 52% left-handed. No significant differences in handedness were found between the bilateral and lateralised groups (t-test, p = 0.232). However, a significant difference in handedness was observed between the typical and atypical groups (t-test, p = 0.002), with the atypical group having significantly lower Edinburgh Handedness Inventory scores, indicating a higher likelihood of left-handedness.

### Average connectivity strength

Following FDR correction for multiple comparisons, only the right supramarginal gyrus (SMG7_R) exhibited a significant association with language lateralisation. Higher FA connectivity strength in this region was observed in typically lateralised individuals (pFDR = 0.016, Cohen’s d = 0.41, Supplementary Figure 1b). No other associations in the FA or rsfMRI modalities remained significant after correction. Additionally, no significant differences were found between lateralised and non-lateralised individuals (Supplementary Figure 1a), and no other nodes showed significant associations in the comparison between typical and atypical lateralisation (Supplementary Figure 1b).

### Structure-function coupling

Overall, the structure-function coupling values were relatively low across all regions (Fisher z < 0.4). Visual inspection of the global structure-function coupling data showed that coupling is highest in amygdala, and lowest in the temporal lobe, based on averaged Fisher z-transformed values (Figure 3). The amygdala (Fisher z = 0.41 on the right, 0.36 on the left) exhibited consistently high coupling values (note that the amygdala is not visible in the figure as it is an internal structure). Similarly, midline regions such as the posterior cingulate gyrus (Fisher z = 0.35 on the left, 0.31 on the right) also demonstrated higher coupling compared to the rest of the nodes. Within cortical regions, coupling was more pronounced in the frontal lobe, where the supplementary motor area (SMA 2, Fisher z = 0.38 on the left, 0.35 on the right) and the pars orbitalis (F3O1, Fisher z = 0.37 on the left, 0.26 on the right) exhibited high values. The temporal lobe showed variable coupling, with regions such as the superior temporal sulcus displaying bilateral coupling (STS_1, Fisher z = 0.32 on both sides), while other temporal areas exhibited comparatively lower values. Coupling appeared relatively symmetric across hemispheres, with bilateral representation in regions such as the amygdala, posterior cingulate gyrus, and superior temporal sulcus. Slightly higher Fisher z-values were observed in the left hemisphere for regions including the supplementary motor area and inferior frontal gyrus. It is important to note, however, that these observations are based on visual inspection of averaged Fisher z-values and do not reflect statistical significance.

**Figure 3.**
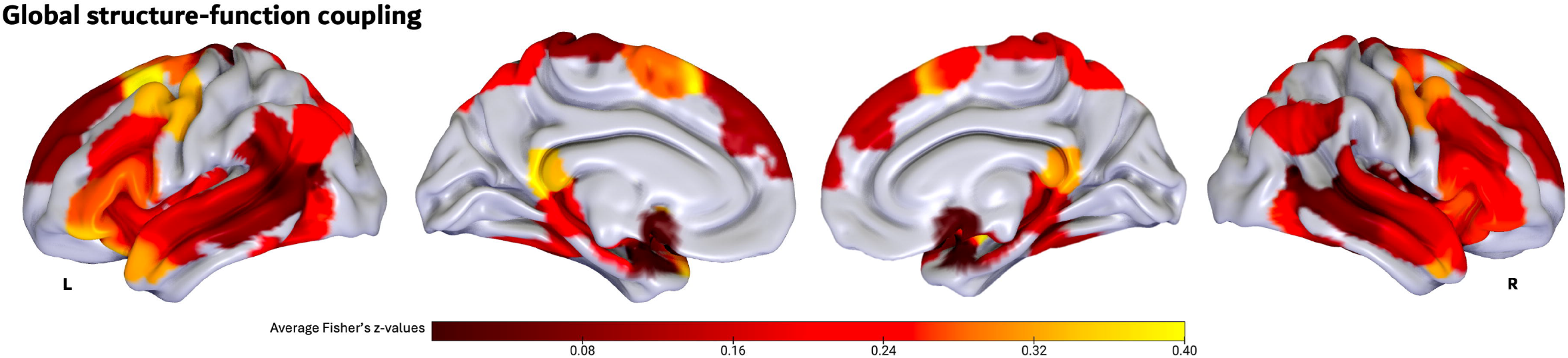
Global structure-function coupling, quantified by average Fisher’s z-values across all participants. The colour map represents the strength of coupling across brain regions, with yellow indicating areas of higher coupling and dark red indicating areas of lower coupling. The lateral, medial, and dorsal views provide a comprehensive visualisation of the spatial distribution of structure-function coupling averaged across the entire cohort.

In the typical versus atypical lateralisation contrast, higher coupling in the typical lateralisation group was observed in the left precentral sulcus (prec3_L) (t = 3.69, Cohen’s d = 0.89, p = 2 × 10^−4^, pFDR = 0.0128), which survived FDR adjustment (Figure 4). Other regions with higher coupling in the typical group, but not surviving FDR adjustment included the right superior temporal sulcus (STS1_R) (t = 2.05, Cohen’s d = 0.5, p = 0.022, pFDR = 0.3888), the left amygdala (AMYG_L) (t = 1.98, Cohen’s d = 0.48, p = 0.024, pFDR = 0.3888), and the left putamen (PUT3_L) (t = 2.02, Cohen’s d = 0.49, p = 0.024, pFDR = 0.3888) (Supplementary Figure 2). Higher coupling in the atypical lateralisation group was observed in the left anterior insula (INSa3_L) (t = 3.14, Cohen’s d = 0.76, p = 7 × 10^−4^, pFDR = 0.0288) and the right middle temporal gyrus (T2_3_R) (t = 3.20, Cohen’s d = 0.77, p = 9 × 10^−4^, pFDR = 0.0288), both of which survived FDR adjustment. Additional regions with higher coupling in the atypical group that did not survive FDR adjustment included the left middle temporal gyrus (T2_3_L) (t = 2.52, Cohen’s d = 0.61, p = 0.0064, q⍰FDR = 0.1365) and the right fusiform gyrus (FUS4_R) (t = 2.05, Cohen’s d = 0.5, p = 0.023, pFDR = 0.2995).

**Figure 4.**
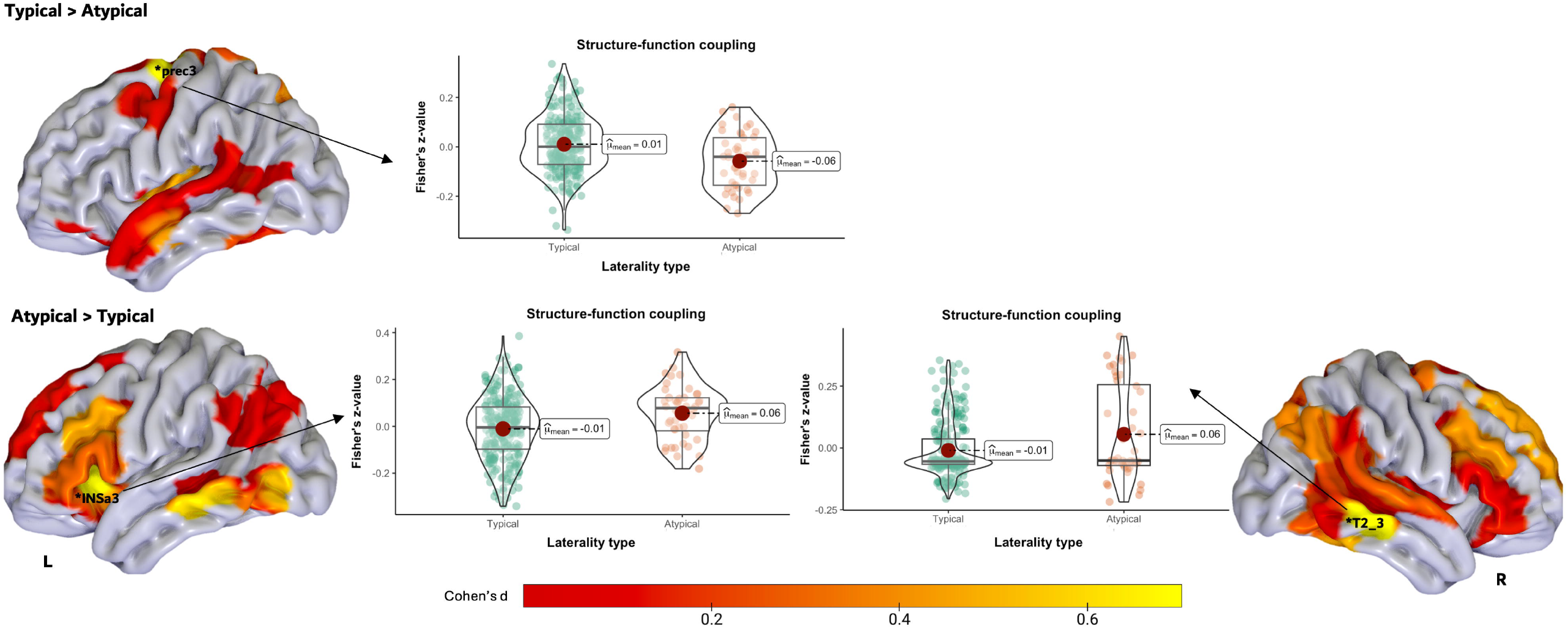
Significant group differences in structure–function coupling between individuals with atypical and typical language lateralisation. This figure presents cortical projections of regions where structure–function coupling significantly differed between individuals with atypical (LI < 18) and typical (LI > 18) language lateralisation for language production. Statistical comparisons were conducted using a t-test within the FSL PALM framework with 10,000 permutations. The colour scale represents Cohen’s d effect sizes, with red indicating smaller effect sizes and yellow indicating larger effect sizes. Regions that survived FDR correction (p_FDR_ < 0.05) are marked with an asterisk (*), highlighting areas where structure–function coupling was either greater in atypical compared to typical individuals or vice versa. Violin plots illustrate the magnitude and direction of effects, showing how coupling strengths differ between groups in regions where atypical lateralisation is associated with either increased or decreased coupling relative to the typical group. Anatomical labels indicate key regions, including the anterior insula (INSa3), upper part of precentral sulcus (prec3), and middle temporal gyrus (T2_3), where significant differences were observed.

In the non-lateralised versus lateralised analysis, higher coupling in people with bilateral language representation was observed in the left anterior insula (INSa3_L) (t = 3.31, Cohen’s d = 0.86, p = 6 × 10^−4^, pFDR = 0.0128), which was the only region to survive FDR adjustment (Figure 5). Additional regions with higher coupling in the bilateral group, but not surviving FDR adjustment, included the right middle temporal gyrus (T2_3_R) (t = 2.63, Cohen’s d = 0.68, p = 0.0072, pFDR= 0.2304), the left middle temporal gyrus (T2_3_L) (t = 2.15, Cohen’s d = 0.56, p = 0.018, pFDR = 0.2832), and the left frontal sulcus (f2_2_L) (t = 1.90, Cohen’s d = 0.49, p = 0.031, pFDR = 0.332) (Supplementary Figure 3). In contrast, regions with higher coupling in the lateralised group included the left precentral sulcus (prec3_L) (t = 3.03, Cohen’s d = 0.79, p = 8 × 10^−4^, pFDR = 0.0512), though it did not survive FDR adjustment, as well as the left superior temporal sulcus (STS2_L) (t = 2.48, Cohen’s d = 0.65, p = 0.0081, pFDR = 0.2592) and the right superior temporal sulcus (STS1_R) (t = 1.85, Cohen’s d = 0.48, p = 0.035, pFDR = 0.7424).

**Figure 5.**
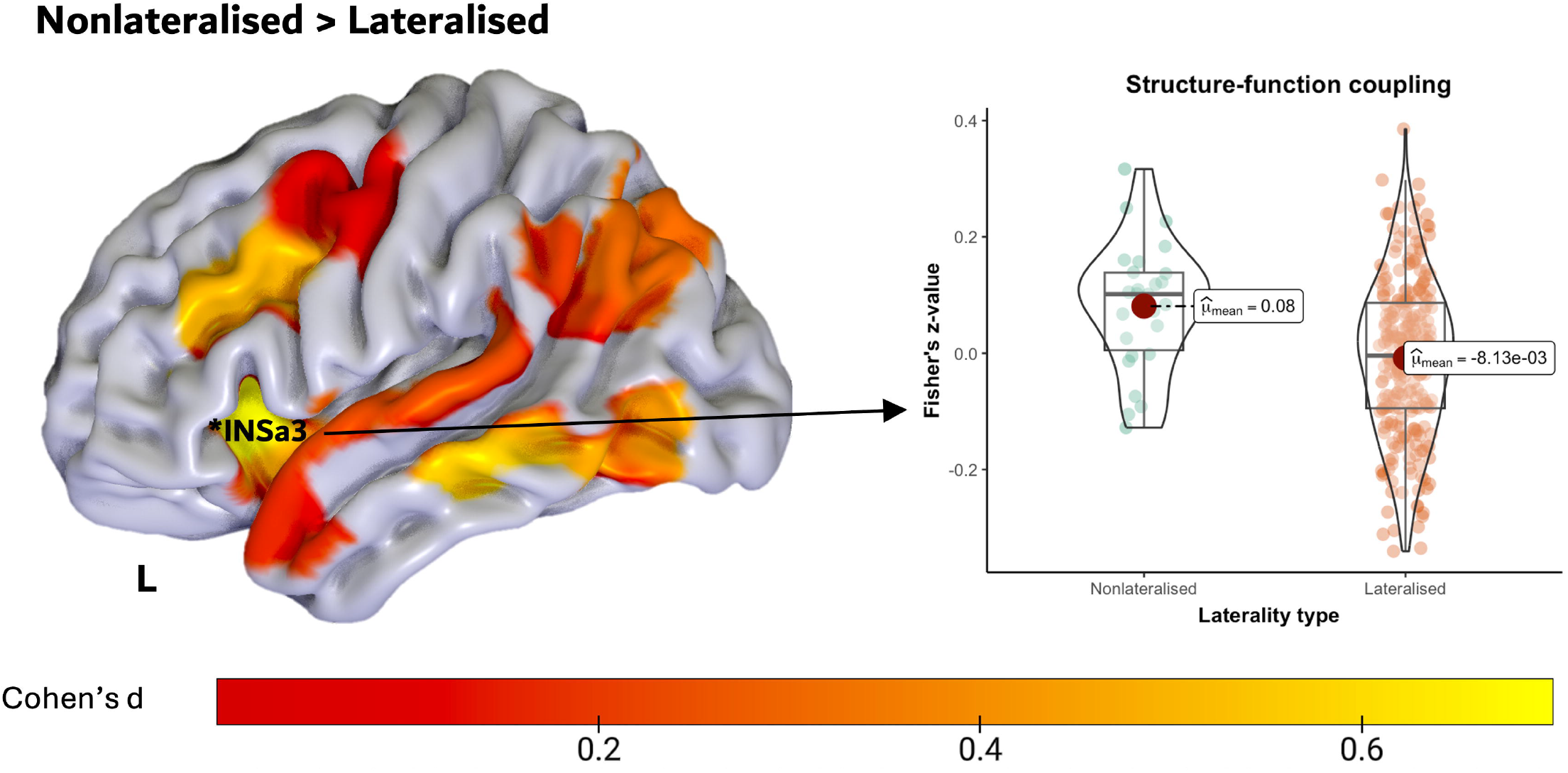
Significant group differences in structure–function coupling between individuals with lateralised and non-lateralised language representation. This figure presents cortical projections of regions where structure–function coupling significantly differed between individuals with lateralised (LI > |20 |) and non-lateralised (20 ≤ LI ≤ 20) language representation for language production. Statistical comparisons were conducted using a t-test within the FSL PALM framework with 10,000 permutations. The colour scale represents Cohen’s d effect sizes, with red indicating smaller effect sizes and yellow indicating larger effect sizes. Regions that survived FDR correction (pFDR < 0.05) are marked with an asterisk (*), highlighting areas where structure–function coupling was significantly greater in non-lateralised compared to lateralised individuals. Violin plots display the distribution of structure– function coupling values for the significant region, with non-corrected p-values reported for visualisation purposes. Anatomical label: INSa3 – anterior insula 3.

### Posthoc analyses

Post-hoc analyses examined structure-function coupling across all connections involving the significant regions: left INSa3, right T2_3, and left prec3. Significant group differences emerged only for connections involving the INSa3_L. Coupling between the left INSa3 and the left INSa2 was significantly higher in atypically lateralised individuals compared to typical individuals (t = 4.12, pFDR = 0.0064, Cohen’s d = 0.71, N = 283) (Figure 6). In lateralised versus non-lateralised individuals, coupling between the INSa3_L and INSa2_L was also higher in the non-lateralised group (t = 5.45, pFDR = 0.00016, Cohen’s d = 1.14, N = 283) (Figure 7). Additionally, a significant difference was observed in the coupling between the INSa3_L and the T2_4_R, with non-lateralised individuals exhibiting higher coupling in non-lateralised individuals (t = 4.04, pFDR = 0.0267, Cohen’s d = 1.24, N = 43).

**Figure 6.**
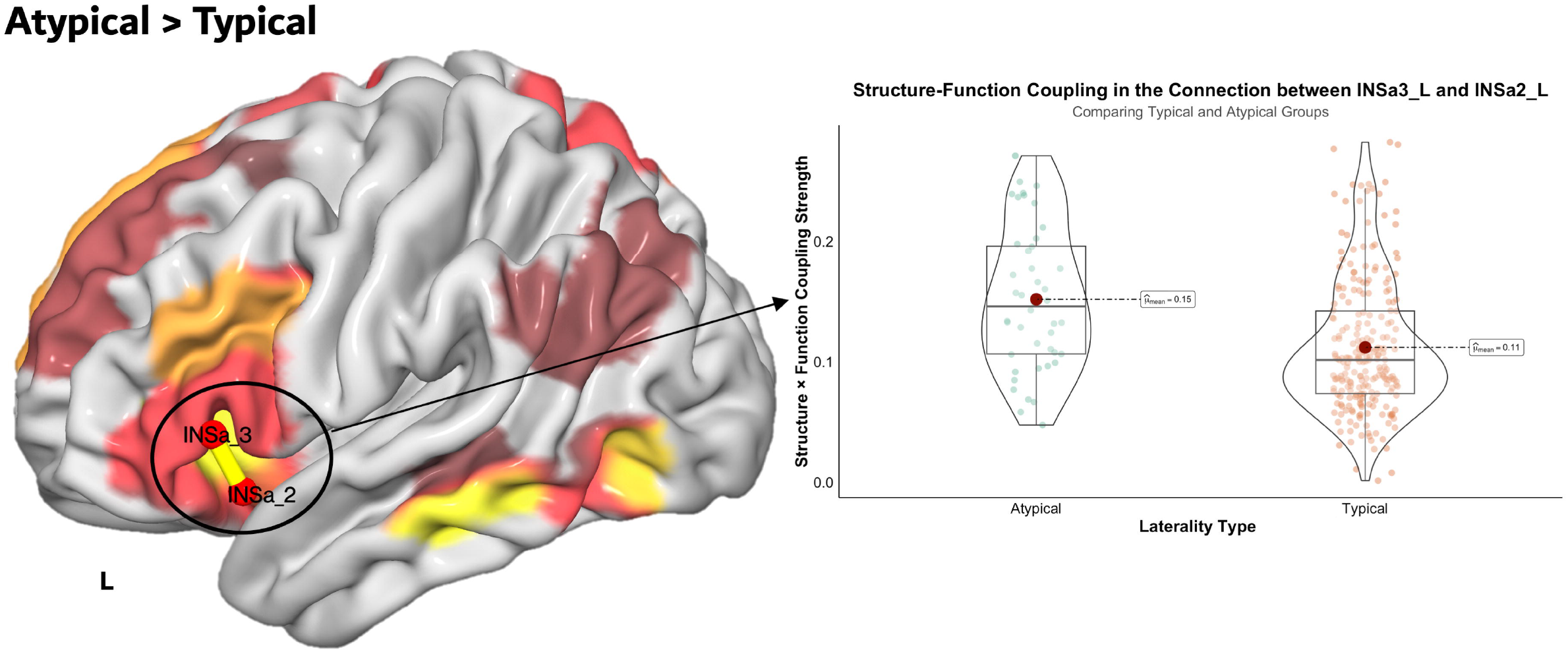
Post-hoc analyses of edge-level structure–function coupling differences between individuals with atypical and typical language lateralisation. This figure presents the connection where post-hoc analyses identified significant group differences in structure–function coupling between atypically (LI < 18) and typically (LI > 18) lateralised individuals. The colours projected on the cortical surface, represents Cohen’s *d* effect sizes as shown in Figure 4. Post-hoc analyses revealed a significant difference in the connection between the left INSa3 and the left INSa2, with atypically lateralised individuals exhibiting significantly higher coupling compared to typically lateralised individuals (*t* = 4.12, *p*FDR = 0.0064, Cohen’s *d* = 0.71, *N* = 283).

**Figure 7.**
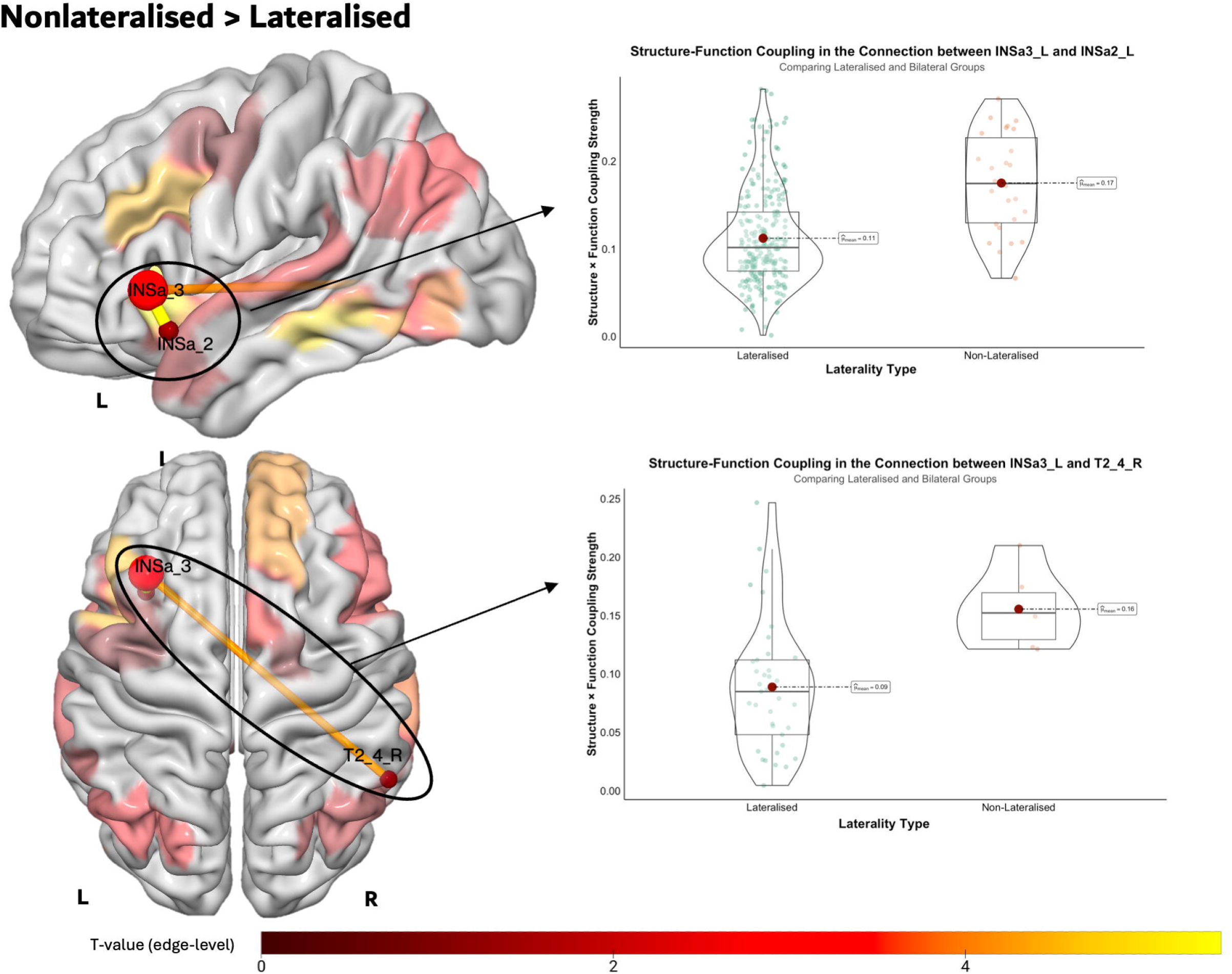
Post-hoc analyses of edge-level structure–function coupling differences between individuals with non-lateralised and lateralised language representation. This figure presents connections where post-hoc analyses identified significant group differences in structure–function coupling between non-lateralised (-20≤ LI ≤ 20) and lateralised (LI > |20|) individuals. The colours projected on the cortical surface represent Cohen’s d effect sizes, as shown in Figure 5, while the colour bar represents t-values for edge-level coupling strength. Non-lateralised individuals exhibited significantly higher coupling between the left INSa3 and the left INSa2 (t = 5.45, pFDR = 0.00016, Cohen’s d = 1.14, N = 283) and between the left INSa3 and the right T2_4 (t = 4.04, pFDR = 0.0267, Cohen’s d = 1.24, N = 43). Violin plots display the distribution of structure–function coupling values for these connections.

No other connections, including those involving T2_3_R or prec3_L, retained significance after multiple comparison correction.

## Discussion

To our knowledge, this is the first study to specifically investigate SC-FC coupling concerning functional language lateralisation. By integrating rsfMRI and diffusion MRI modalities, we explored nodal connectivity strength and SC-FC coupling to uncover distinct patterns across language laterality groups. Consistent with our hypotheses, SC-FC coupling differentiated typical and atypical individuals. Unexpectedly, however, atypical individuals were not marked by reduced coupling, as hypothesised; instead, they exhibited heightened coupling in specific regions, including the left anterior insula and right middle temporal gyrus, while SC-FC couplingof the left precentral sulcus was associated with typical language dominance. Non-lateralised individuals showed increased coupling in the left anterior insula compared to lateralised groups, suggesting its role in supporting non-lateralised language networks. Analyses of rsfMRI and diffusion MRI FA connectivity independently revealed no significant group differences in rsfMRI, while FA connectivity strength in the right supramarginal gyrus was associated with typical language dominance.

### SC-FC coupling – biological relevance

SC-FC coupling reflects the interplay between structural and functional connectivity, offering insights into how brain anatomy supports functional dynamics. Our findings reveal that SC-FC differentiates typical and atypical language lateralisation groups, with distinct patterns emerging in the left superior precentral sulcus, left anterior insula, and right MTG. High SC-FC coupling is associated with neural efficiency, where functional activity is tightly constrained by structural pathways, enabling energy-efficient and predictable communication (Haier et al., 1988; Neubauer & Fink, 2009). This efficiency stems from structural scaffolding, as strong coupling reinforces stable, anatomically grounded networks (Buckner & Krienen, 2013). For instance, the strong coupling in typical lateralisation likely reflects reliance on well-established language networks that support consistent hemispheric dominance for language tasks.

At a global level, overall coupling values were low, consistent with previous studies showing reduced coupling in higher-order regions (Figure 3; Buckner & Krienen, 2013; Chamberland et al., 2017; Vázquez-Rodríguez et al., 2019). This trend aligns with the atlas used, which focuses on supramodal regions involved in language sentence production (Labache et al., 2019). Lower SC-FC coupling in higher-order regions, such as those involved in language, reflects their greater cognitive flexibility and capacity for dynamic reconfiguration (Chamberland et al., 2017; Smith, 2016; Zimmermann et al., 2018). This flexibility reduces reliance on structural pathways, enabling adaptive functional networks for complex linguistic and cognitive tasks (Diamond, 2020).

In contrast, regions distinguishing laterality groups exhibited stronger coupling, reflecting signalling constraints essential for polysensory integration (Buckner & Krienen, 2013; Vázquez-Rodríguez et al., 2019). These regions likely achieve efficient language processing through stable SC-FC coupling, avoiding frequent reconfiguration or heavy neuromodulation (Suárez et al., 2020). This pattern aligns with the Neural Efficiency Hypothesis of Intelligence, which posits that individuals with higher cognitive abilities exhibit more efficient neural processing, requiring less activation during cognitive tasks (Haier et al., 1988; Popp et al., 2024; Wang, 2021). While our study did not study language performance differences between groups, heightened SC-FC coupling in higher-order regions, such as the inferior frontal lobe, has been linked to improved reading performance in other studies (Kotoski et al., 2024), suggesting a similar mechanism may support language proficiency.

Contrary to expectations, atypical individuals did not exhibit reduced SC-FC coupling. Variability in SC-FC coupling across regions and groups reinforces the idea that coupling is highly context-dependent and not uniformly distributed. For example, while typical individuals showed increased coupling in the left precentral sulcus, atypical individuals did not show a mirrored increase in the right precentral sulcus (Figure 4). This lack of mirroring suggests that atypical individuals do not simply replicate typical lateralisation patterns in the opposite hemisphere. Instead, they exhibit distinct SC-FC coupling organisation, supporting the idea that typical and atypical individuals represent two distinct populations with different structural and functional language organisation. The observed patterns align with the concept of the brain operating near criticality, where regions balance stability and adaptability to optimise processing efficiency (Park & Friston, 2013). This balance enables different lateralisation profiles to rely on distinct structural-functional configurations, underscoring the diversity in neural mechanisms underlying language dominance.

### Topology of SC-FC coupling in relation to language lateralisation

Our study identified the SC-FC coupling in the upper part of the left precentral sulcus was associated with typical language lateralisation. The prominence of this region in our findings likely reflects its unique developmental, structural, and functional characteristics. Emerging around the 24th week of gestation and running approximately parallel to the central sulcus (Chi et al., 1977), the precentral sulcus (PrCS) provides early anatomical scaffolding essential for articulatory and phonological processing (Matchin & Hickok, 2020). The morphology of PrCS demonstrates a consistent left-hemispheric bias, defined by a higher incidence of its presence in the left hemisphere compared to the right, across both humans and non-human primates (Juch et al., 2005; S. S. Keller et al., 2007; Ono et al., 1990). Deeper left PrCS morphology has been linked to right-handedness (Amunts et al., 1996; Foundas et al., 1998; Klöppel et al., 2007; Willems & Hagoort, 2009), which, in turn, correlates with left-hemisphere language dominance (Johnstone et al., 2021; Knecht et al., 2000). Keller et al. (2007, 2009) further demonstrated that sulci neighbouring the inferior frontal gyrus (IFG) (such as the PrCS and diagonal sulcus) contribute to leftward asymmetry in the IFG by increasing intrasulcal area, potentially facilitating leftward language lateralisation (Powell et al., 2012). Given that our findings specifically pertain to SC-FC coupling in the upper PrCS, caution is warranted in directly applying broader conclusions from whole-PrCS morphology. Further studies incorporating detailed morphological analyses of PrCS subdivisions could clarify its precise contribution to language lateralisation.

The anterior insula was a key region where heightened SC-FC coupling differentiated atypical from typical language lateralisation, as well as non-lateralised individuals from lateralised ones. This finding was not replicated in separate analyses of rsfMRI or diffusion MRI FA connectivity, highlighting the unique value of SC-FC coupling in capturing the integrated dynamics of structural and functional networks underlying language lateralisation. While leftward structural asymmetry of the anterior insula has been linked to typical language dominance (Biduła & Króliczak, 2015; S. S. Keller et al., 2011), atypical individuals often show reduced or reversed asymmetry (Greve et al., 2013). In our study, however, heightened SC-FC coupling in the left anterior insula was associated with atypical lateralisation, raising questions about its left-lateralised focus. Network-based studies have previously shown associations between FA connectivity in both the left and right anterior insula and strongly atypical lateralisation (Andrulyte, Zago, et al., 2025), though these findings reflect a different scope – FA connectivity rather than SC-FC coupling. Conceptualised as a “code translator,” the anterior insula is thought to bridge structural and functional networks, enabling efficient communication between neural systems that support linguistic and non-linguistic processes, including attention and salience detection (Cai & Van Der Haegen, 2015; Molnar-Szakacs & Uddin, 2022; Uddin, 2015). In people with atypical and non-lateralised language representation, heightened coupling may indicate a supportive mechanism that allows for enhanced cross-hemispheric communication while remaining structurally anchored within left-lateralised pathways. This integrative role may explain why the anterior insula remains persistently involved in atypical and non-lateralised individuals, functioning as an adaptive mediator of language networks that supports the dynamic coordination of structural and functional asymmetries.

A key distinction emerged between right-lateralised and bilateral individuals in how SC-FC coupling was organised within the anterior insula. In atypical individuals (including right-lateralised and non-lateralised), SC-FC coupling was primarily driven by intrahemispheric connectivity within the left anterior insula – specifically, between INSa_3 and INSa_2 (Figure 6). In non-lateralised as compared to lateralised individuals, an additional coupling effect was observed in the interhemispheric connectivity between the left INSa_3 and the right middle temporal gyrus (MTG; T2_4) (Figure 7). This suggests that while both right-lateralised and bilateral individuals rely on intrahemispheric insular connectivity within the left hemisphere, supported by structural scaffolding (Biduła & Króliczak, 2015), non-lateralised individuals additionally recruit interhemispheric connections involving the MTG. The functional significance of this additional connectivity remains unclear, but its selective presence in these individuals suggests it may reflect an alternative strategy for language network organisation in non-lateralised individuals. Further research is needed to determine whether this connectivity pattern has functional consequences or represents variability in SC-FC coupling across individuals.

Interestingly, while our results indicated increased SC-FC coupling in the right MTG (T2_3) in atypically lateralised individuals, post-hoc analyses did not identify significant edge-level coupling differences between groups. This raises questions regarding the precise role of right MTG connectivity in atypical lateralisation. Previous studies have demonstrated that increased FA nodal degree in the right MTG is associated with right hemisphere language dominance (Zahnert et al., 2023) and semantic performance in patients with left temporal lobe epilepsy (Audrain et al., 2018). These findings suggest that the right MTG may act as a compensatory structure for linguistic functions in the absence of typical leftward dominance. While only the right MTG survived multiple comparisons in our analysis, the left MTG also showed a trend toward significance (pFDR = 0.1365, Cohen’s d = 0.5; Supplementary Figure 2), suggesting that both hemispheres may contribute to language processing in atypically lateralised individuals. This aligns with previous network-based diffusion MRI studies, which have demonstrated that individuals with atypical language dominance exhibit increased connectivity between the MTG and other temporal regions, both inter- and intra-hemispherically (Andrulyte, Zago, et al., 2025).

### Methodological considerations

The methodological framework of this study reflects both its strengths and areas requiring further exploration. One of the key methodological considerations was focusing exclusively on positive rsfMRI correlations for SC-FC coupling, prioritising clearer interpretability of results. Although negative correlations between nodes in rsfMRI studies are documented, especially in higher-order regions (Fox et al., 2005; Uddin et al., 2009), their biological significance remains debated (J. B. Keller et al., 2015; Murphy et al., 2009; Nir et al., 2020; Weissenbacher et al., 2009). Although this choice may limit the scope of interpretation, focusing on positive correlations aligns with the majority of SC-FC coupling studies and supports robust and interpretable analyses (Baum et al., 2020).

Interpreting the structural connectivity underlying SC-FC coupling also presents methodological challenges, particularly in relation to FA. While FA is widely employed in diffusion MRI studies, it does not directly reflect white matter integrity but rather serves as a proxy for microstructural organisation (Jones et al., 2013; Rheault et al., 2020). Its sensitivity to fibre architecture, particularly in regions with multiple crossing fibres, can obscure meaningful biological interpretations (Jeurissen et al., 2013). In association and commissural pathways, where multiple fibre populations intersect, the diffusion tensor model is unable to resolve distinct orientations, leading to artificially reduced FA values even when connectivity remains intact (Tournier et al., 2011). As a result, variations in FA between groups may arise from diverse sources – including differences in fibre complexity, neuroinflammatory processes, or acquisition-related artefacts – rather than reflecting true differences in connectivity (Figley et al., 2022). To mitigate these limitations, future studies should consider integrating more advanced diffusion models that refine white matter characterisation (Daducci et al., 2015; Mayer et al., 2022; H. Zhang et al., 2012).

Another important limitation relates to the diffusion MRI FA connectivity matrices, where 70–95% of connections were missing for key nodes, including the left anterior insula, left precentral sulcus, and right MTG. This highlights that SC-FC coupling in these regions may reflect only a subset of connections rather than a comprehensive view of node connectivity. Additionally, the use of a group atlas assumes uniform functional boundaries across individuals, which may fail to account for significant interindividual variability, particularly in transmodal cortices where functional boundaries are known to differ (Gordon et al., 2017). This potential misalignment could affect the accuracy of SC-FC coupling measurements, leading to certain regions being underrepresented or overemphasised in the analysis (Suárez et al., 2020).

Future research should address these limitations by investigating interindividual variability through participant-specific parcellations to better capture the functional organisation of higher-order regions. Examining the connections to and from specific regions, such as the anterior insula, MTG, and precentral sulcus, will be critical for understanding their roles in hemispheric language dominance. Additionally, studies should employ a range of language tasks and alternative atlases to provide a more comprehensive understanding of the functional and structural dynamics underlying SC-FC coupling. These steps are vital to advancing our understanding of SC-FC coupling’s role in language lateralisation and delineating the specific contributions of key cortical regions.

## Conclusion

Our findings establish SC-FC coupling as a valuable approach for distinguishing typical and atypical language lateralisation and lateralised and non-lateralised profiles. The left anterior insula emerged as a key region, supporting left-hemisphere access in atypical and non-lateralised individuals, while the right MTG was uniquely linked to atypical lateralisation, potentially aiding cross-hemispheric processing. Typical language dominance was linked to stronger coupling in the left superior precentral sulcus, reaffirming its critical role in hemispheric asymmetry. These results emphasise the importance of these regions in language lateralisation and invite further exploration into their specific contributions.

## Funding

Ieva Andrulyte – BBSRC DTP training grant BB/T008695/1

## CREDIT authorship contribution statement

Ieva Andrulyte – conceptualisation, data analysis, funding acquisition, writing – original draft

Laure Zago – data collection, writing – review & editing Gael Jobard – data collection, methodology

Herve Lemaitre – data collection Peter N Taylor – methodology

Francois Rheault – methodology, writing – review & editing

Marc Joliot – data collection, methodology, writing – review

Laurent Petit – data collection, methodology, writing – review & editing, supervision

Simon S Keller – conceptualisation, writing – review & editing, supervision

## Competing Interests

Authors declare no competing interests

## Data availability statement

This study is based on the BIL&GIN dataset, which is accessible through collaborative research agreements designed to foster scientific cooperation. Details on data access policies can be found in Mazoyer et al. (2016). While the dataset itself is not publicly available, all analysis scripts used in this study are openly accessible on GitHub at https://github.com/andrulyte/SF-Coupling.

---

**Supplementary Figure 1.**
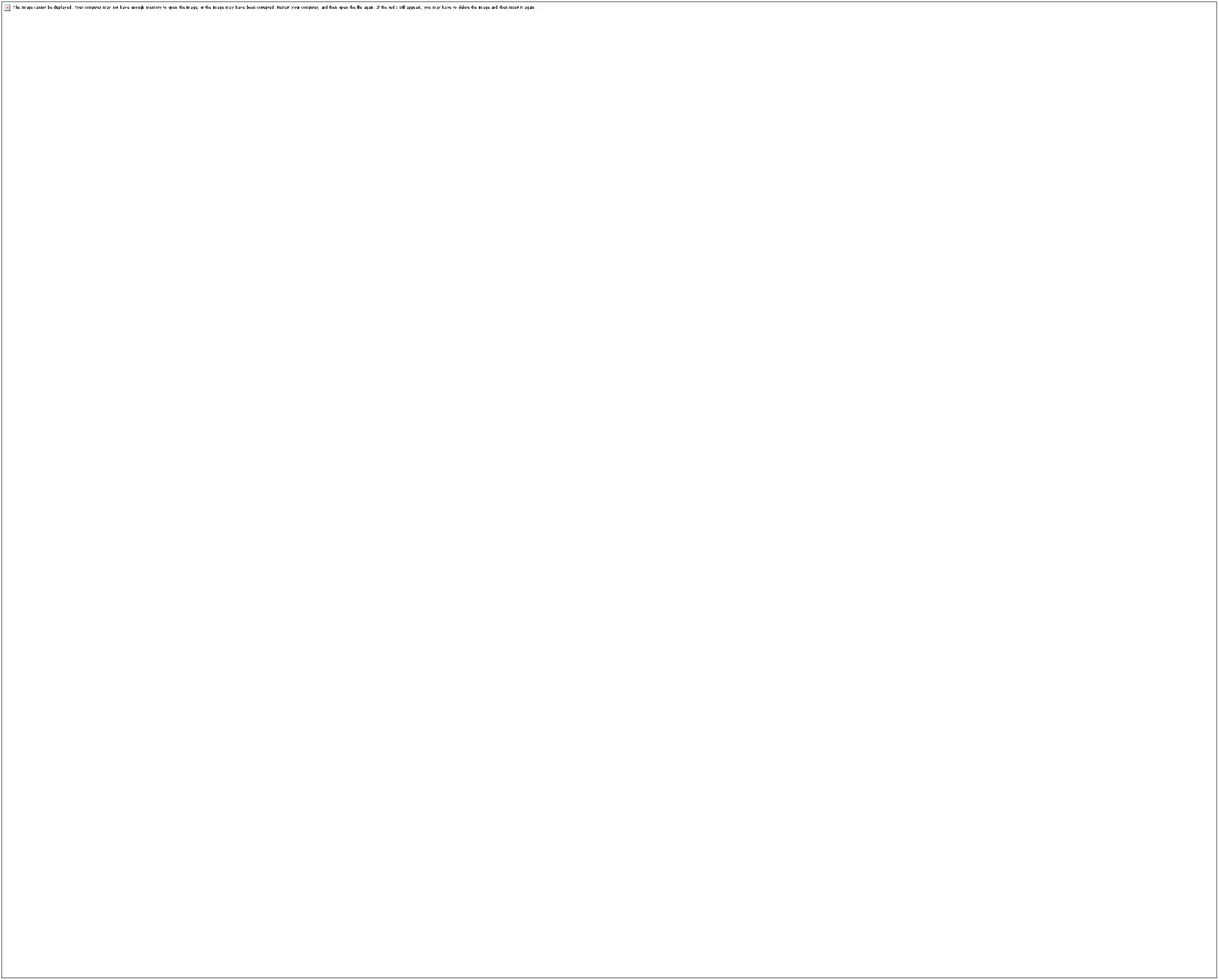
Group-averaged connectivity strength for FA and rsfMRI data, visualised using Cohen’s d. This figure examines connectivity related to language production, specifically comparing groups based on their degree of language lateralisation. Panel (a) compares individuals with lateralised language representation (left- or right-hemisphere dominance for language) to those with non-lateralised (or bilateral) language representation. Panel (b) compares individuals with typical left-hemisphere lateralisation for language to those with atypical lateralisation (either right-hemisphere dominance or bilateral representation). Blue regions indicate greater connectivity strength in the non-lateralised group compared to the lateralised group (a) or in the atypically lateralised group compared to the typically lateralised group (b). Conversely, red regions indicate greater connectivity strength in the lateralised group compared to the non-lateralised group (a) or in the typically lateralised group compared to the atypically lateralised group (b). Only in the comparison between typically and atypically lateralised groups (panel b) did a region survive FDR correction (pFDR < 0.05): *SMG7 - supramarginal gyrus 7, which showed greater connectivity in the typically lateralised group.

**Supplementary Figure S2.**
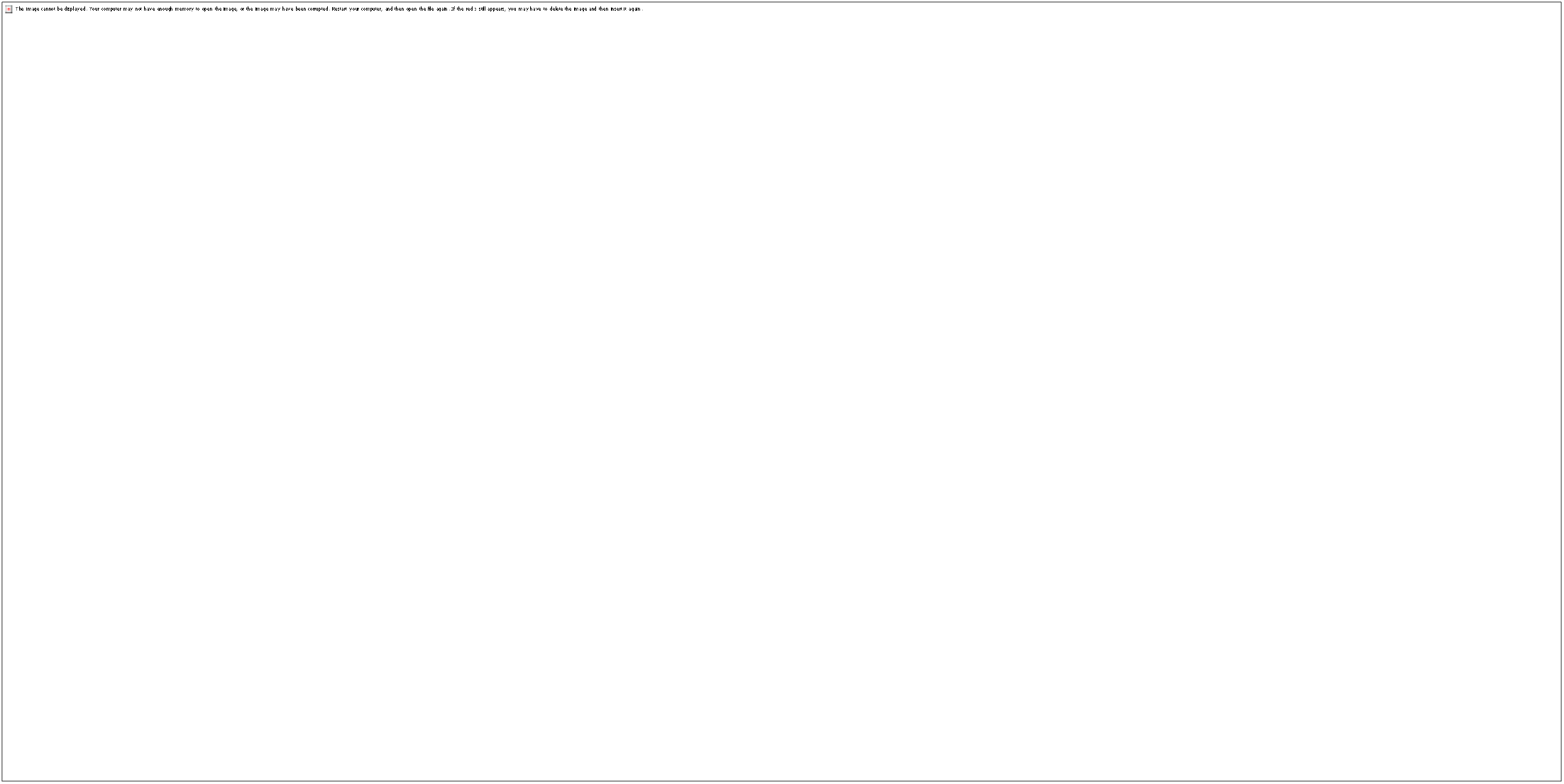
Whole-brain differences in structure–function coupling between typical and atypical language lateralisation groups. This figure presents whole-brain projections of structure–function coupling differences between individuals with typical (LI > 18) and atypical (LI < 18) language lateralisation. Results were obtained using a t-test within the FSL PALM framework with 10,000 permutations. The colour scale represents Cohen’s d effect sizes. Regions that survived FDR correction (pFDR < 0.05) are marked with an asterisk (*), as shown in Figure 4.

**Supplementary Figure S3.**
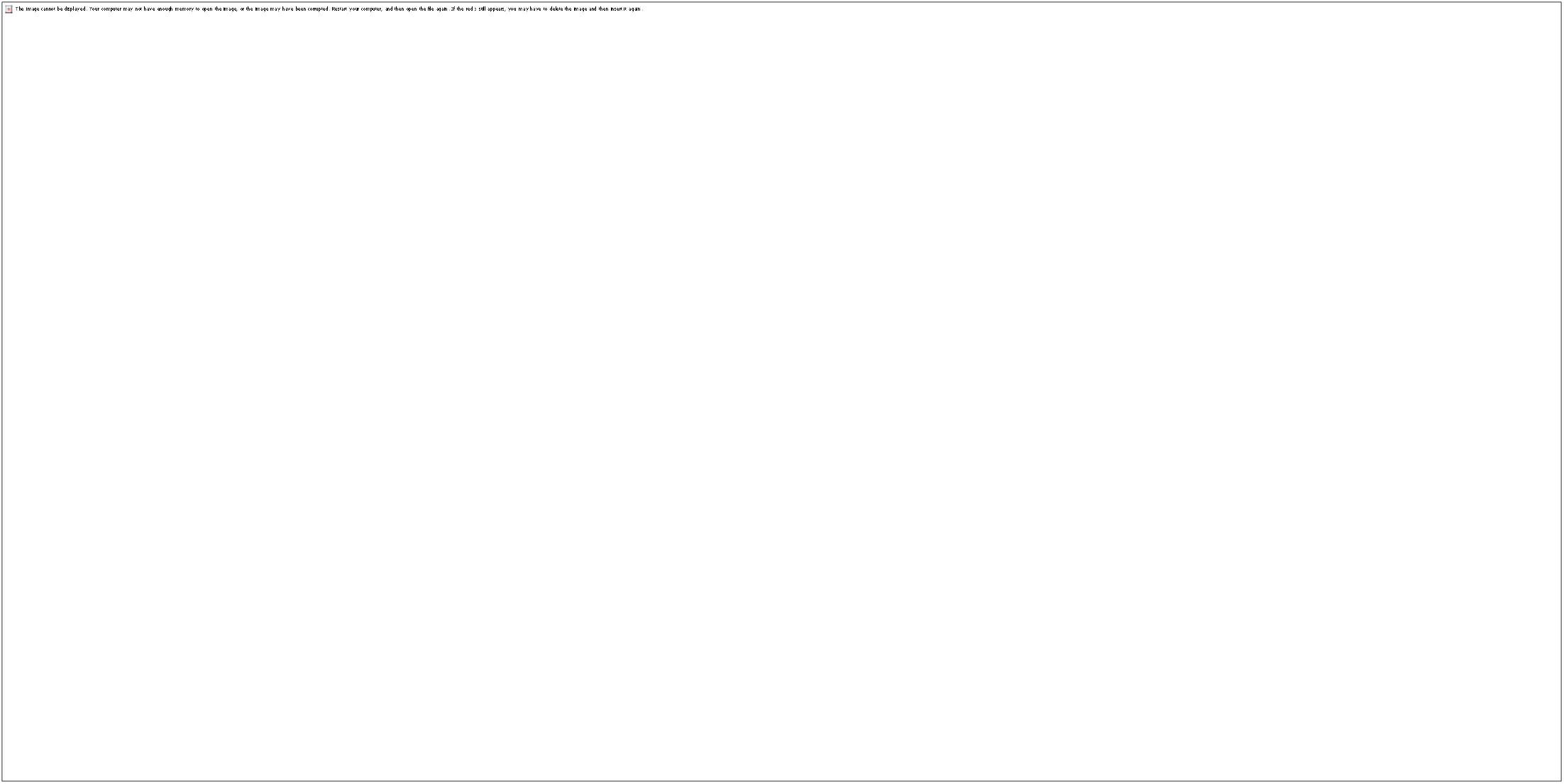
Whole-brain differences in structure–function coupling between individuals with lateralised and bilateral language representation. This figure presents whole-brain projections of structure–function coupling differences between individuals with lateralised (LI > |20 |, encompassing both left (LI > 20) and right (LI < 20) lateralisation) and bilateral (20≤ LI ≤ 20) language representation during language production. Statistical comparisons were conducted using a t-test within the FSL PALM framework with 10,000 permutations. The colour scale represents Cohen’s d effect sizes, with red indicating smaller effect sizes and yellow indicating larger effect sizes. Regions that survived FDR correction (pFDR < 0.05) are marked with an asterisk (*).

## Notes

### Competing Interest Statement

The authors have declared no competing interest.

### Summary of Updates

Updating the text, including CREDIT, Github link, and updating figures

