## Supplementary figures and images for "Brain structure-function coupling – relationship with language lateralisation"

### Supplementary Figure 1

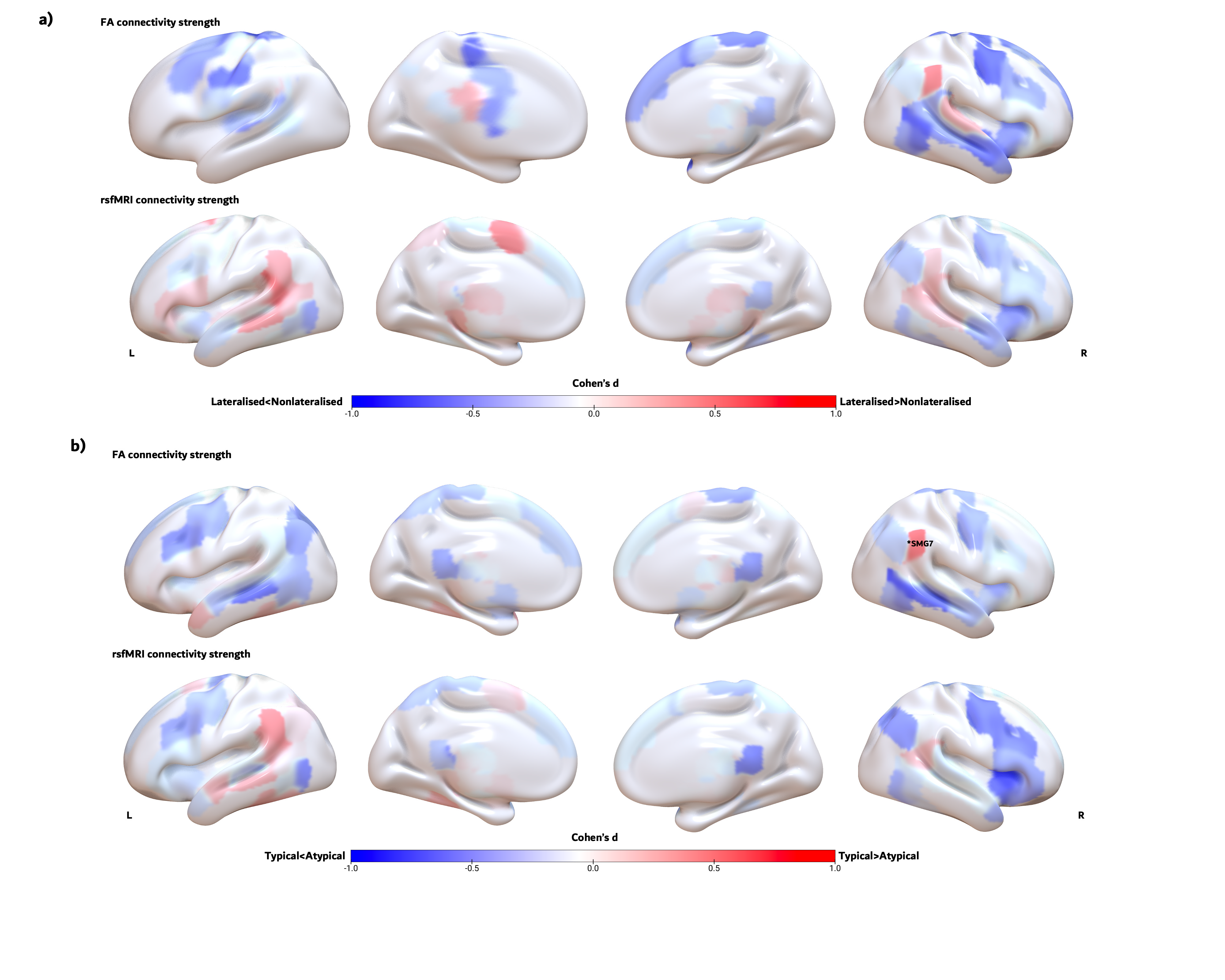

### Supplementary Figure 2

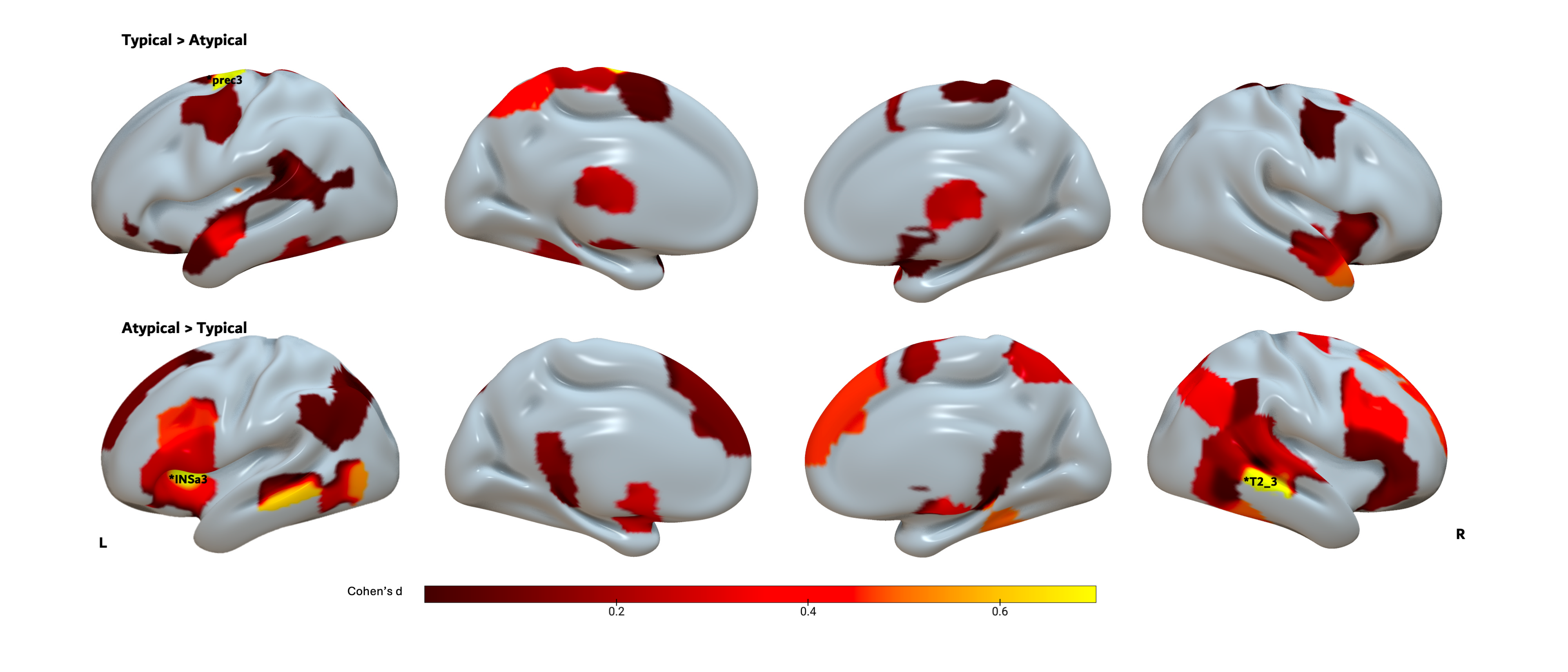

### Supplementary Figure 3

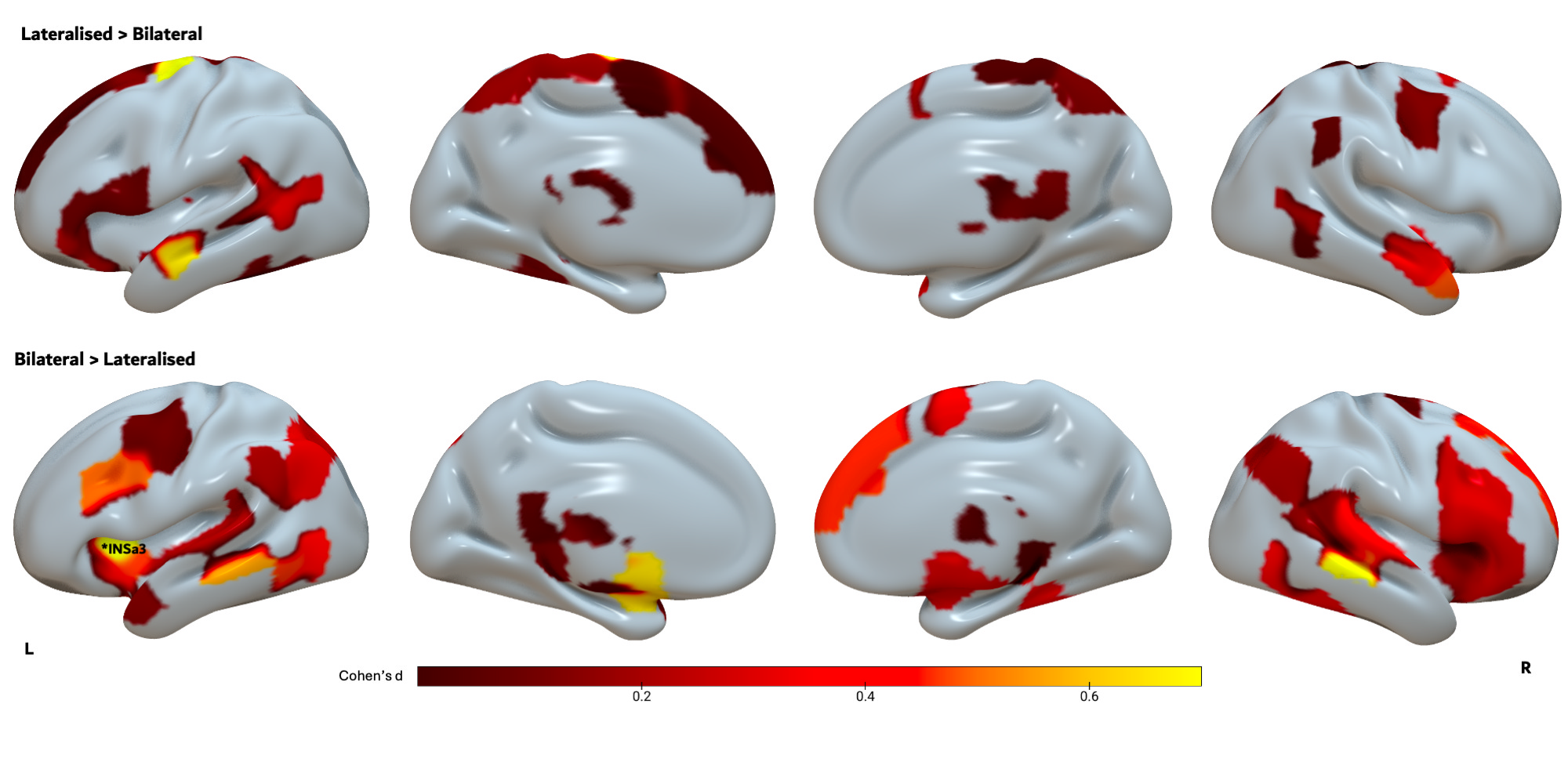
